# The chikungunya virus E1 glycoprotein fusion loop and hinge alter glycoprotein dynamics leading to cell and host specific changes in infectivity

**DOI:** 10.1101/2023.11.03.565585

**Authors:** Sara A. Thannickal, Leandro Battini, Sophie N. Spector, Maria G. Noval, Diego E. Álvarez, Kenneth A. Stapleford

## Abstract

Alphaviruses infect both mammals and insects, yet the distinct mechanisms that alphaviruses use to infect different hosts are not well defined. In this study, we characterize CHIKV E1 variants in the fusion loop (E1-M88L) and hinge region (E1-N20Y) *in vitro* and *in vivo* to understand how these regions of the E1 glycoprotein contribute to host-specific infection. Through cell culture assays, we found that CHIKV E1-N20Y enhanced infectivity in mosquito cells while the CHIKV E1-M88L variant enhanced virus binding and infectivity in both BHK-21 and C6/36 cells, and led to changes in the virus cholesterol-dependence in BHK-21 cells. Given these *in vitro* results and that residue E1-M88L is in a defined Mxra8 interacting domain, we hypothesized that this residue may be important for receptor usage. However, while the CHIKV E1-M88L variant increased replication in Mxra8-deficient mice compared to WT CHIKV, it was attenuated *in vitro* in mouse fibroblasts, suggesting that residue E1-M88 may function in a cell-type dependent manner to alter entry. Finally, using molecular dynamics to understand how potential changes in the E1 glycoprotein may impact the CHIKV glycoprotein E1-E2 complex, we found that E1-M88L and other E1 domain II variants lead to changes in both E1 and E2 dynamics. Taken together, these studies show that key residues in the CHIKV E1 fusion loop and hinge region function through changes in E1-E2 dynamics to facilitate cell- and host-dependent entry.

**Importance:** Arthropod-borne viruses (arboviruses) are significant global public health threats, and their continued emergence around the world highlights the need to understand how these viruses replicate at the molecular level. The alphavirus class II glycoproteins are critical for virus entry in mosquitoes and mammals, yet how these proteins function is not completely understood. Therefore, to address these gaps in our knowledge, it is critical to dissect how distinct glycoprotein domains function *in vitro* and *in vivo*. Here, we show that changes in the CHIKV E1 fusion loop and hinge contribute to host-specific entry and E1-E2 dynamics, furthering our knowledge of how alphaviruses infect mammals and insects.

## Introduction

Alphaviruses are single-stranded positive-sense RNA viruses responsible for causing arthritic and encephalitic diseases worldwide (1) (2) (3). This genus includes arthropod-borne viruses (arboviruses) such as eastern equine encephalitis virus, chikungunya virus (CHIKV), and Mayaro virus. As there are no effective antiviral therapeutics available, understanding the mechanisms of how alphaviruses replicate and enter cells is crucial for preventing and controlling future epidemics.

CHIKV is a re-emerging alphavirus spread by *Aedes (Ae.)* mosquitoes. CHIKV causes disease consisting of rash, arthralgia, and in severe cases can cause neurological and cardiac manifestations (4) (5) (6) (7) (8). Importantly, CHIKV continues to emerge with a significant increase in CHIKV cases in South America in 2023, emphasizing the importance of further studying the mechanisms, transmission, and spread of CHIKV (9) (10). The alphavirus genome consists of two open reading frames, the first encoding 4 nonstructural proteins (nsP1—nsP4) and the latter encoding the structural proteins, including capsid, E3, 6K/TF, and the class II fusion glycoproteins E1 and E2 (11). Embedded in the virion membrane, the CHIKV E1 and E2 glycoproteins are essential in the early stages of receptor-mediated endocytosis. The E1-E2 heterodimer is disrupted as the endosomal acidic pH sparks a conformational change where E1 trimerizes and further facilitates the fusion of viral and endosomal membranes (12) (13) (14) (15). The alphavirus E1 glycoprotein is comprised of 3 domains (E1-I, E1-II, and E1-III), where domain II contains the fusion loop, *ij* loop, and *bc* loop that have been previously studied for their importance in attachment, fusion and infectivity (16) (17) (18). Notably, during a significant outbreak in 2005 on La Réunion Island, an E1-A226V mutation in the domain II *ij* loop was identified and found to increase fitness in *Ae. albopictus* mosquitoes (10) (19).

In previous studies, we have shown that the CHIKV E1 glycoprotein β-strand c plays an important role in the entry, infectivity, fusion, and dissemination of CHIKV both *in vitro* and *in vivo* (20). In these studies, we generated a panel of amino acid substitutions in the β-strand c residue V80 and identified a network of second-site mutations between V80 and multiple domains of the E1 glycoprotein using *in vivo* and *in vitro* evolution. Specifically, we identified two novel second-site E1 mutations that were found in *Ae. aegypti* mosquitoes infected with the CHIKV E1 variant V80Q. These mutations (E1-M88L and E1-N20Y) were initially identified by an increased plaque size, and were hypothesized to play important roles in the stability of the E1-V80Q variant as well as potentially for the evolution and transmission of CHIKV.

In this study, we generated the CHIKV E1-M88L and N20Y variants and used both *in vitro* and *in vivo* approaches to characterize how these residues contribute to CHIKV entry and pathogenesis. We found that CHIKV E1-M88L and E1-N20Y significantly changed the host- and cell-type infection of CHIKV *in vitro*. However, CHIKV E1-M88L had no phenotype differences in mosquitoes or C57BL/6J mice. When evaluating usage of the CHIKV receptor Mxra8, the E1-M88L variant showed increased viral titers in Mxra8-deficient mice, suggesting it may be using alternative routes of infection compared to wild-type CHIKV. Finally, using molecular simulations to understand how CHIKV E1 domain II variants may impact the E1-E2 dimer, we found that changes in the E1 fusion loop, β-strand c, and *ij* loop can alter the dynamics of both E1 and E2, indicating changes in E1 can have significant impacts on the entire glycoprotein spike. Taken together, these studies highlight how changes in the E1 fusion loop can impact virus binding and infection and how changes in the E1 protein contribute to multiple steps in the entry process beyond fusion.

## Results

### CHIKV E1-M88L and E1-N20Y enhance infection in a host-specific manner

In a previous study to understand how the CHIKV E1 glycoprotein β-strand c contributes to infection, we mutated the CHIKV E1 β-strand c residue V80 to every possible amino acid and identified a variant of V80 (E1-V80Q) that was characterized by a small plaque phenotype and significant attenuation *in vitro* and *in vivo* (20). When we infected *Ae. aegypti* mosquitoes with CHIKV E1-V80Q, we observed two emerging second-site E1 glycoprotein variants associated with increased plaque size. One variant, E1-M88L, is located in the conserved E1 fusion loop of domain II and the other, E1-N20Y, is located in the E1 hinge region of domain I (**Fig. 1A**). Given the observation of an increased plaque size, we hypothesized that the emergence of E1-M88L and/or E1-N20Y was rescuing E1-V80Q through genetic interactions. Unfortunately, when we attempted to generate the E1-V80Q single and double variants to address this question, we found that the E1-V80Q single variant was genetically unstable, reverting to the wild-type valine or obtaining the E1-M88L mutation. Therefore, without a proper control we could not explore interactions between E1-V80Q and E1-M88L or E1-N20Y. Nonetheless, given the emergence of E1-M88L with E1-V80Q and the location of these residues in the E1 glycoprotein, we hypothesized that M88 and N20 are important residues for virus entry and pathogenesis.

**Figure 1:**
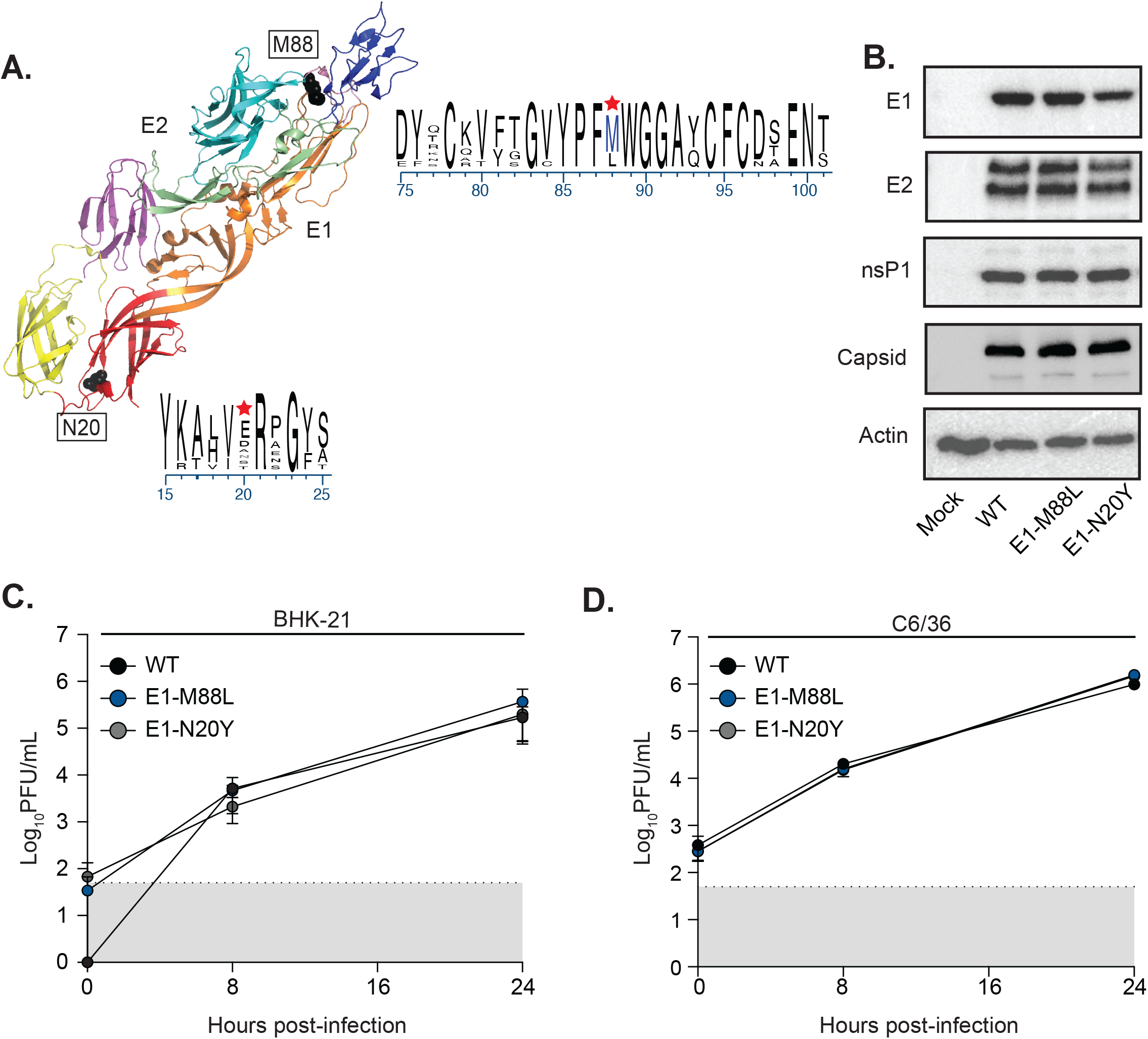
CHIKV E1-M88L and E1-N20Y variant conservation and growth kinetics. (**A**) PyMol structure (PDB: 3N42) depicting the CHIKV E1 and E2 glycoprotein. Domains are color-coded as follows: E1-1 (red), E1-II (orange), Fusion Loop (pink), E1-III (yellow), E2-A (light blue), E2-B (dark blue), E2-C (purple), E2-β ribbon (green). The E1-M88 and E1-N20 positions are in black and labeled. A composite logo of the fusion loop and part of the E1-1 domain alignments across 12 alphaviruses are shown, with a red star indicating the CHIKV E1-M88 and E1-N20 position. (**B**) BHK-21 cells were electroporated with *in vitro* transcribed RNA of each virus or mock transfected. Cells were lysed 48 hours post electroporation and CHIKV E1, E2, nsP1, capsid, and actin accumulation was visualized by western blotting. Blots are a representation from at least three independent trials. BHK-21 cells (**C**) and C6/36 cells (**D**) were infected at an MOI of 0.1 for 1 hour, washed with PBS, and supernatants were collected at the indicated timepoints. Viral titers were quantified via plaque assay. The dotted line and gray shaded area represent the limit of detection (LOD). Data represent three independent trials, and no statistical significance was found via two-way ANOVA. The average and standard error of the mean (SEM) are shown for all data.

To characterize how these residues contribute to CHIKV infection, we first aligned both the fusion loop and part of the domain I hinge region across multiple alphaviruses. We found that E1-M88 is conserved in 75% of the alphaviruses of our alignment (**Fig. 1A**). Interestingly, the leucine variant is found in Aura virus (AURV), Middelburg virus (MIDV) and Una virus (UNAV), suggesting this residue may provide some advantage in other viruses. On the contrary, E1-N20 is highly variable among alphaviruses (**Fig. 1B**). To begin to characterize these variants *in vitro*, we first electroporated mammalian BHK-21 cells with each viral *in vitro* transcribed (IVT) RNA and lysed the cells after 48 hours to assess protein accumulation by western blot. We found that there were no differences in protein accumulation between wild-type (WT) CHIKV and either E1 variant (**Fig. 1B**). Furthermore, we also looked at the growth kinetics of each mutant in both mammalian BHK-21 (**Fig. 1C**) and insect C6/36 *Aedes* (*Ae*.) *albopictus* (**Fig. 1D**) cells, yet found no growth difference between the viruses.

However, given that the CHIKV E1-V80 residue contributes to CHIKV infectivity and E1-M88L is found in the fusion loop, we hypothesized that these variants contribute to virus infectivity. To test this hypothesis, we performed an ammonium chloride bypass assay. We adsorbed virus to BHK-21 and C6/36 cells for 1 hour at 4°C and then incubated the cells at 37°C and treated with 20 mM ammonium chloride at different time points to neutralize the endosomal and lysosomal pH and block virus infection and spread (**Fig. 2A and B**). We found that E1-M88L led to faster infection and enhanced CHIKV infectivity in BHK-21 cells while E1-N20Y behaved like WT CHIKV (**Fig. 2A**). When we completed the same experiment with insect cells, we found that there was minimal infection in C6/36 cells until the 60 minutes post-infection timepoint, at which we see an increased infectivity for both E1-M88L and E1-N20Y (**Fig. 2B**). These results suggest that the E1-M88L variant increases infection in both hosts while E1-N20Y is insect cell specific. To investigate if this increased infectivity was due to cell binding, we incubated both mammalian and insect cells with each virus for 30 mins at 4°C in the presence of 20 mM ammonium chloride and quantified the number of membrane-bound virus via qPCR (**Fig. 2C**). We found that E1-M88L increased binding in both cell types, while E1-N20Y bound to cells similar to WT CHIKV. Lastly, as CHIKV entry is cholesterol-dependent, we sought to determine if there were any differences in cholesterol-dependent entry in the E1 variants. We depleted BHK-21 cells of cholesterol using varying concentrations of methyl-beta-cyclodextrin (MβCD), infected cells with each virus, then added 20 mM ammonium chloride to block spread and therefore determine the entry efficacy of each virus. We found that E1-M88L showed a decrease in cholesterol dependency, as the virus was still able to infect cells at higher concentrations of MβCD compared to E1-N20Y and WT (**Fig. 2D**). These results show that while E1-N20Y had an increased infectivity in C6/36 cells, E1-M88L is able to bind, fuse, and enter both mammalian and insect cells more efficiently compared to WT, and therefore we chose to focus on the E1-M88L variant for the remainder of this study.

**Figure 2:**
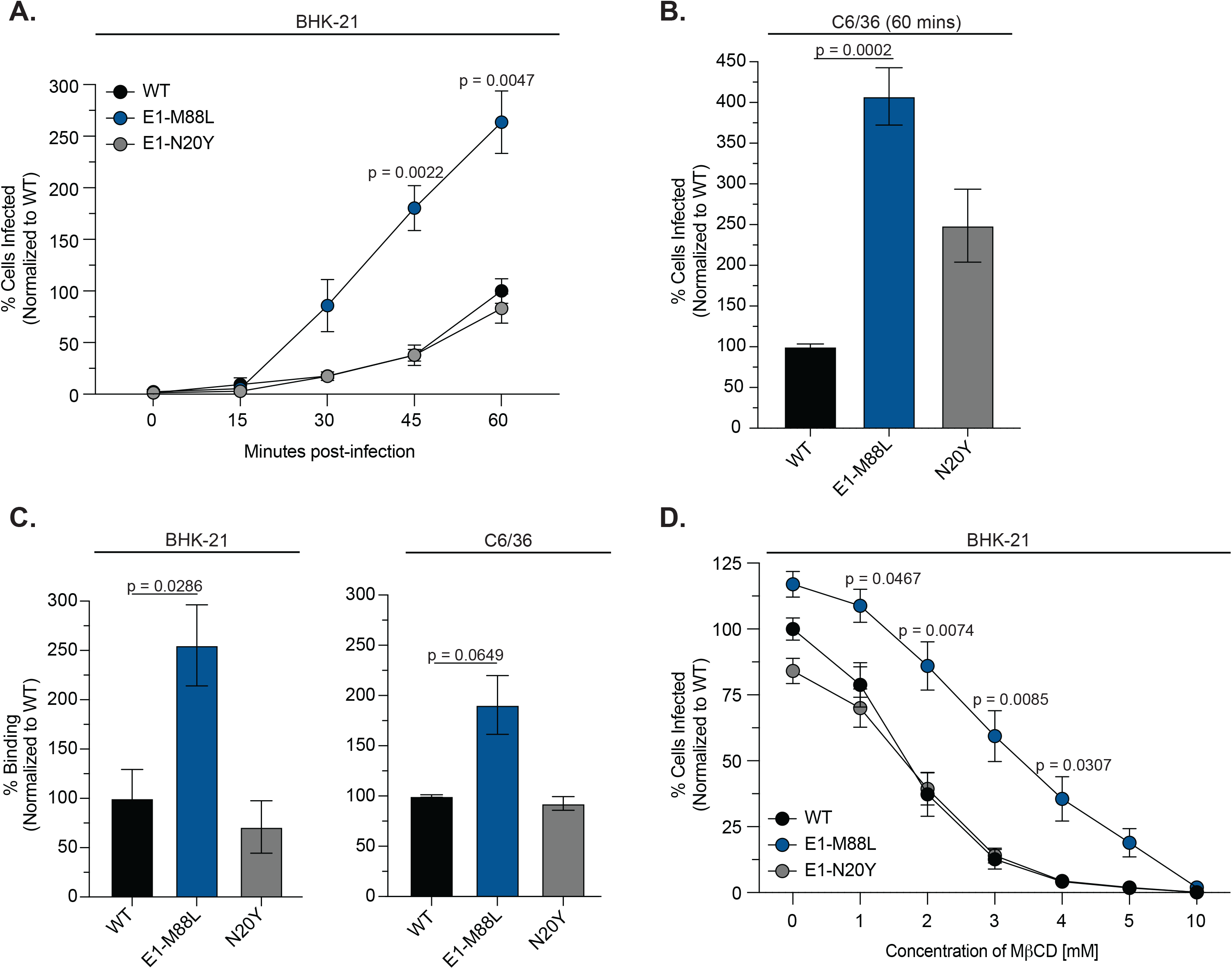
CHIKV E1-M88L and E1-N20Y influence binding and infectivity in a host-dependent manner. (**A**) BHK-21 cells were infected with WT CHIKV or each E1 variant expressing Zs-Green at an MOI of 1 and treated with 20 mM ammonium chloride at the indicated timepoints post-infection. Cells were fixed and stained with DAPI 24 hours post infection. Infected cells quantified using a CX7 high-content microscope. Data represent three independent trials with internal duplicates. Statistical significance was found by two-way ANOVA and indicated with p-values shown. (**B**) C6/36 cells were infected with WT CHIKV or each E1 variant expressing ZsGreen at an MOI of 0.1 for 1 hour and then treated with 20 mM ammonium chloride. Cells were fixed and stained at 24 hours post infection and infected cells quantified as above. Data represent four independent trials with internal duplicates. Statistical significance was found by Kruskal-Wallis one-way ANOVA test and indicated with p-values shown. (**C**) Binding assays were performed on BHK-21 and C6/36 cells. Cells were incubated on ice with each virus at an MOI of 100 (based on viral genomes) for 1 hour. After a cold PBS wash, virus-bound cells were collected and RNA genomes were quantified using qPCR. Data represent at least three independent experiments, with internal duplicates. A Mann-Whitney test was used to determine statistical significance, and indicated p-values are shown. (**D**) BHK-21 cells were pre-treated with methyl-beta-cyclodextrin (MßCD) for 1 hour, wash once with PBS, and cholesterol-depleted cells were infected with each CHIKV virus at an MOI of 1 for 1 hour before adding 20 mM ammonium chloride. Cells were fixed and stained 24 hours post infection and quantified as above. Data represent three independent trials with internal duplicates. Statistical significance was found by two-way ANOVA and indicated with p-values shown. The average and SEM are shown for all data.

### CHIKV E1-M88L does not alter RNA replication but does increase extracellular virus

Given the dramatic enhancement of infectivity of the CHIKV E1-M88L variant, we found it interesting there were no differences in growth kinetics between E1-M88L and WT in cell culture. One hypothesis for these results may be that while the CHIKV E1-M88L particle is more infectious, there is a defect in another step in the life cycle, such as RNA replication. To test this hypothesis, we introduced the E1-M88L variant into a virus expressing a Firefly luciferase reporter that is expressed under active replication through a subgenomic promoter (**Fig. 3A**). We transfected BHK-21 cells with each CHIKV *in vitro* transcribed RNA and measured intracellular luciferase activity at 4, 6, 8, and 24 hours post-transfection (**Fig. 3A and B**). Prior to the luciferase readout, we transferred the culture supernatant to naïve cells and measured the intracellular luciferase activity 24 hours later in order to evaluate infectious particle production (**Fig. 3A and C**). While there were no differences in luciferase activity between WT CHIKV and E1-M88L (**Fig. 3B**), we did observe an increase in luciferase activity from the supernatant at 8 hours post transfection compared to WT CHIKV (**Fig. 3C**). These results indicate that while there are no differences in subgenomic replication, CHIKV E1-M88L may be producing infectious particles faster than WT or more infectious particles, which may lead to enhanced infection.

**Figure 3:**
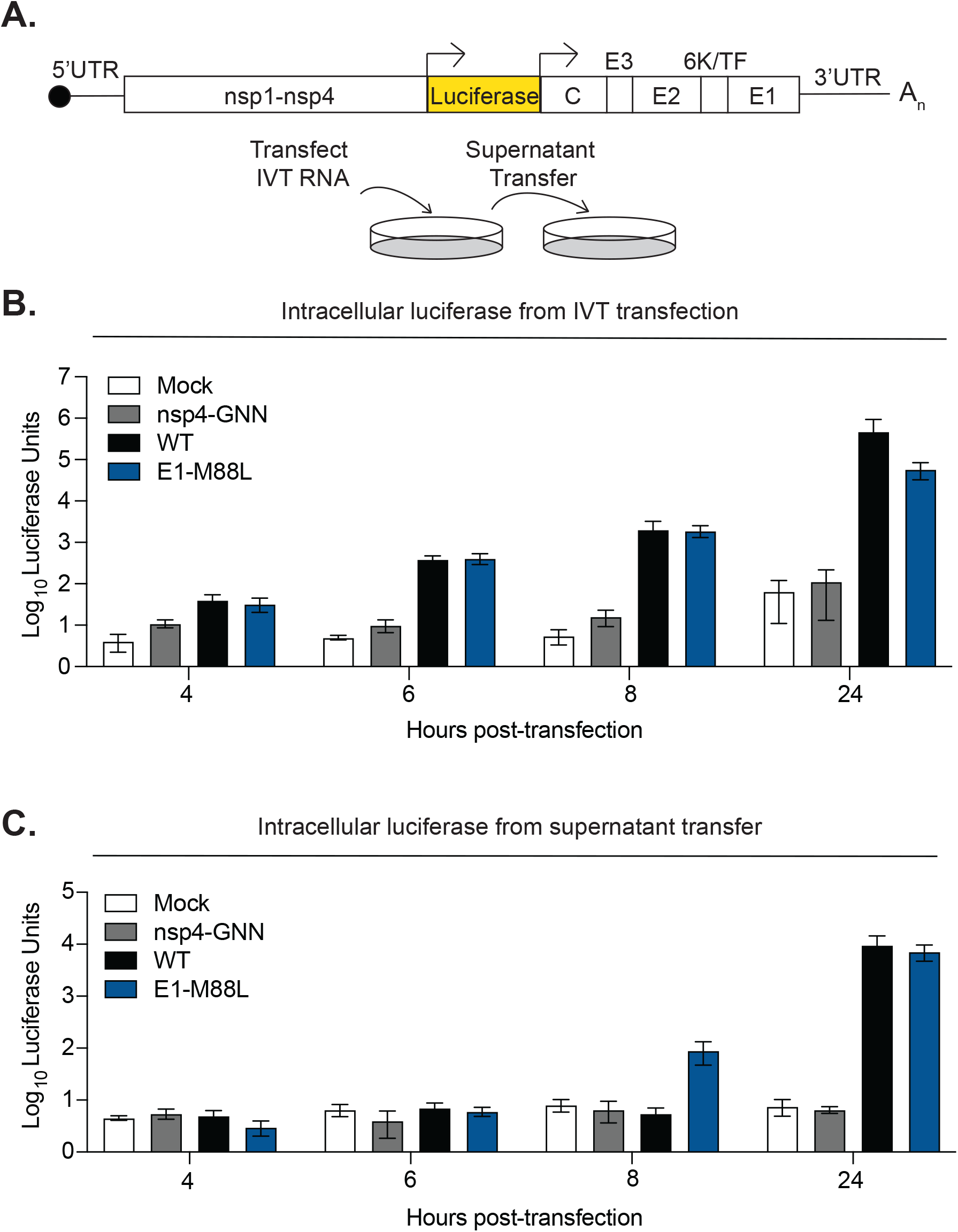
WT CHIKV and E1-M88L subgenomic replication and particle production. (**A**) A schematic of the CHIKV genome expressing the firefly luciferase reporter (top) and schematic of the experiment (bottom). (**B**) BHK-21 cells were transfected with WT CHIKV, nsp4-GNN or E1-M88L firefly IVT RNAs using Lipofectamine 2000. At each indicated timepoint, the supernatant was transferred to naïve BHK-21 cells and the cells lysed to quantify luciferase activity. (**C**) 24 hours post infection of the naïve BHK-21 cells, the cells were lysed and luciferase activity quantified. Data represent four independent trials. No statistical significance was found via two-way ANOVA. The average and SEM are shown for all data.

### The CHIKV E1-M88L variant shows no major advantage in mosquitoes or wild-type mice

As CHIKV E1-M88L was found as a second-site mutation from a CHIKV E1-V80Q infection in *Ae. aegypti* mosquitoes and E1-M88L enhanced infectivity and binding in C6/36 cells, we wanted to see if there were differences in infection and dissemination of CHIKV E1-M88L in a mosquito model. We infected *Ae. aegypti* mosquitoes with 10^6^ PFU of WT or CHIKV E1-M88L, collected the mosquito bodies and legs and wings after 7 days post-infection, and quantified infectious virus by plaque assay. We found no differences in infection or dissemination to the legs and wings between WT CHIKV and E1-M88L (**Fig. 4A and 4B**). In addition, since CHIKV E1-M88L enhanced infectivity and binding in BHK-21 cells, we hypothesized this residue may play a role in infection in mice. We infected C57BL/6J mice with 1000 PFU of each virus via the footpad and harvested footpad, ipsilateral calf and quadricep muscle, and serum at 2, 3, and 5 days post-infection to quantify viral particles by plaque assay (**Fig. 4C-4E**). We found that CHIKV E1-M88L had no significant advantage in mice at most timepoints with the exception of a slight decrease in infectious particles at 3 dpi in the footpad and serum. These results suggest that the function of the CHIKV E1-M88L variant plays cell-specific roles that could be masked in whole organism models.

**Figure 4:**
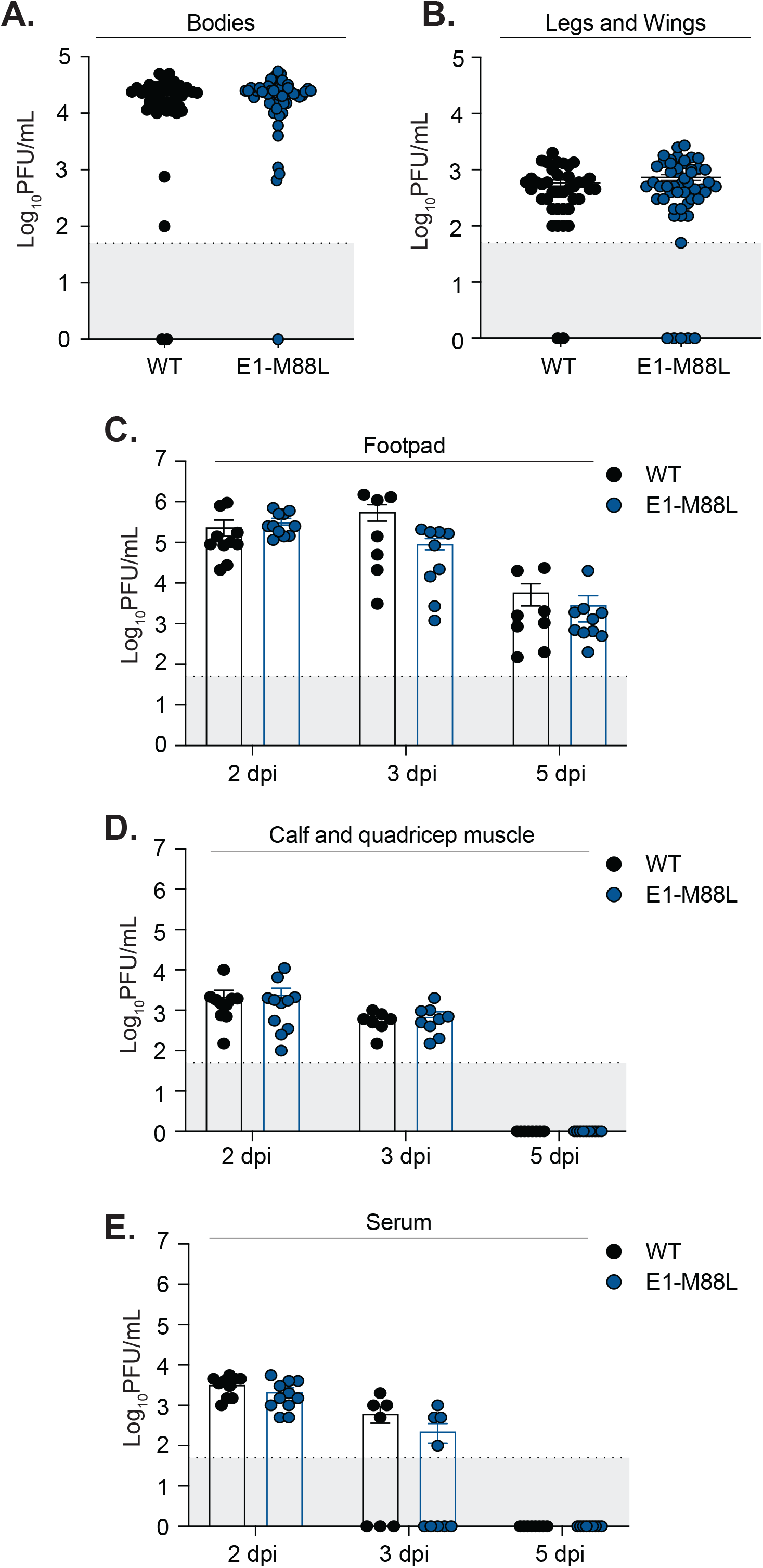
WT CHIKV and E1-M88L replication in *Aedes aegypti* mosquitoes and C57BL/6J mice. *Aedes aegypti* mosquitoes were fed a viral blood meal containing 10^6^ PFU/mL of WT CHIKV or E1-M88L virus. Mosquitoes were maintained for 7 days post-infection and viral titers were quantified via plaque assay in bodies (**A**) or legs and wings (**B**). Data represent two independent infectious with at least n = 43 mosquitoes. Male and female 5 to 7-week-old C57BL/6J mice were infected with 1000 PFU of WT CHIKV or E1-M88L virus via footpad injection. Mice were euthanized at the corresponding timepoint and the footpad (**C**), the ipsilateral calf and quadricep muscle (**D**) and serum (**E**) harvested. Infectious titers were determined via plaque assay. Data represent at least 2 independent infections with at least n = 9 mice. The dotted line and gray shaded area represent the limit of detection (LOD). No statistical significance was found via Mann-Whitney test. The average and SEM are shown for all data.

### CHIKV E1-M88L increases replication at the site of infection in Mxra8-deficient mice, but not in Mxra8-deficient mouse fibroblasts *in vitro*

The Matrix Remodeling Associated 8 (Mxra8) protein is an important receptor for CHIKV and other arthritogenic alphaviruses (2) (21) (22). CHIKV residue E1-M88 is in proximity to the Mxra8 receptor interaction domain (23), suggesting its ability to potentially alter binding to the CHIKV E1-E2 dimer and impact infection (**Fig. 5A**). To investigate this hypothesis, we infected Mxra8-deficient and WT litter mate control mice with 1000 PFU of either WT CHIKV or E1-M88L virus via footpad injection. At 3 days post-infection, we harvested the footpad and ipsilateral calf and quadricep muscle and quantified infectious virus by plaque assay. We found that while WT and E1-M88L replicated to the same levels in the footpad of wild-type mice, the E1-M88L variant lead to a statistically significant increase in infectious particles in the footpad of Mxra8-deficient mice (**Fig. 5B and 5C**). However, this phenotype was not seen in the ipsilateral calf and quadricep muscle. These results suggest that the CHIKV E1-M88L variant may be less dependent on Mxra8 and/or may use an alternative route to bind, enter and infect *in vivo*.

**Figure 5:**
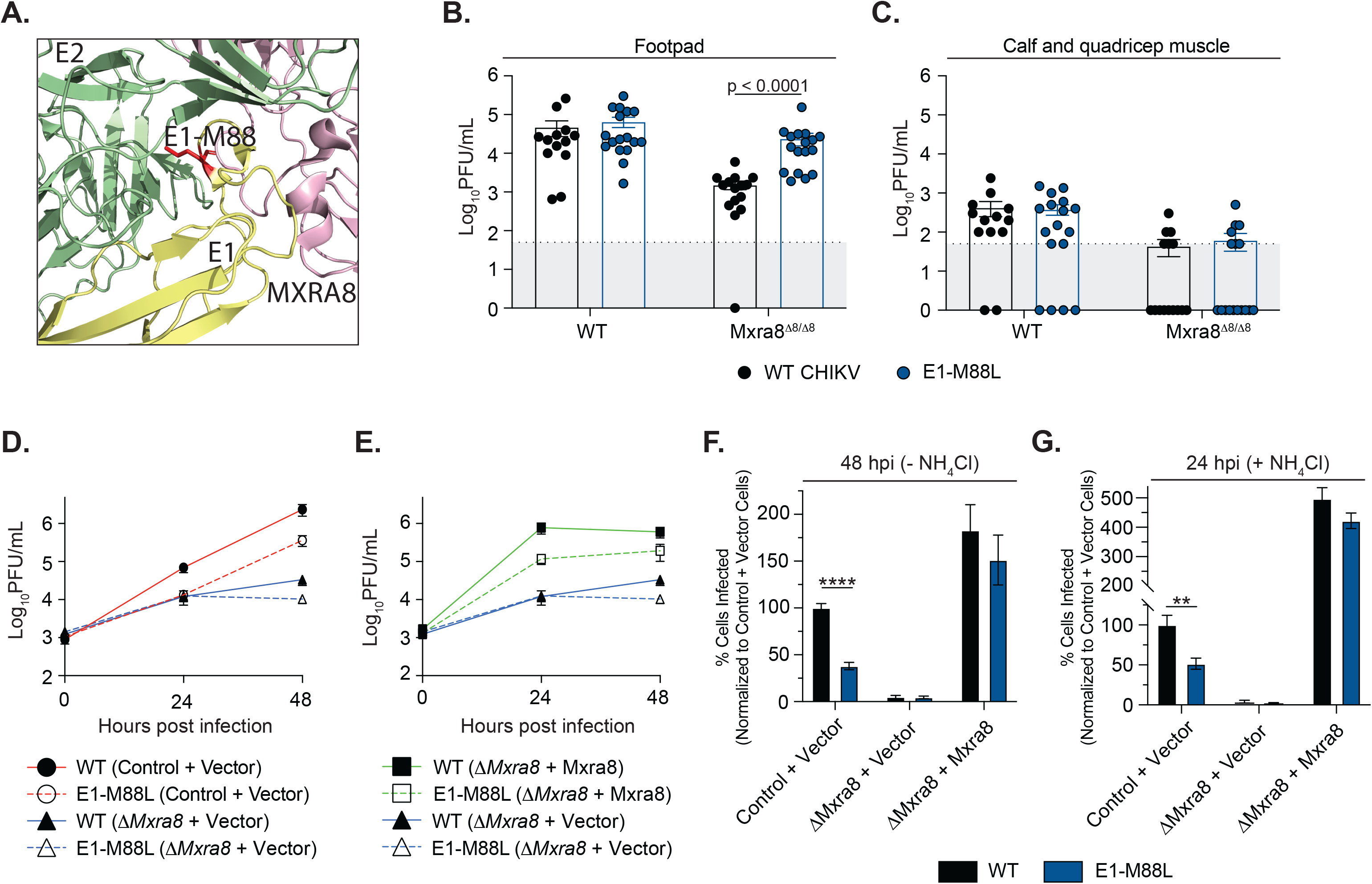
CHIKV E1-M88L replicates in Mxra8-deficient mice but is attenuated in mouse fibroblasts. (**A**) Crystal structure of the CHIKV E1 glycoprotein in yellow, E2 in green, Mxra8 receptor in pink, and the E1-M88 residue in red. Male and female 5–7-week-old WT or *Mxra8^Δ8/Δ8^* mice were infected with 1000 PFU of WT CHIKV or E1-M88L virus via footpad injection. Mice were euthanized at 3 days post-infection, and the footpad (**B**) and ipsilateral calf and quadricep muscle (**C**) harvested to quantify infectious titers via plaque assay. Data represent at least three independent infectious with at least n =13. The dotted line and gray shaded area represent the limit of detection (LOD). (**D** and **E**) Virus growth in wild-type NIH 3T3 mouse embryonic fibroblasts (MEF), Mxra8-deficient MEFs, or MEFs expressing Mxra8 in trans. NIH 3T3 MEF cells were infected with each virus expressing Zs-Green with an MOI of 5 for 1 hour, washed with PBS, and complete media added. Supernatants were collected at the indicated timepoints and infectious titers were quantified via plaque assay. At 48 hpi cells were fixed, stained with DAPI, and the number of infected cells quantified as above (**F).** (**G**) Each cell line was incubated with each virus at an MOI of 5 for 1 hour followed by the addition of 20 mM ammonium chloride. At 24 hpi, cells were fixed and stained for DAPI and infected cells were quantified using a CX7 high-content microscope. Data represent at least two independent trials in triplicate. Multiple Mann-Whitney tests were performed with p-values representing **p < 0.01 and ****p < 0.0001. The average and SEM are shown for all data.

Given the results in the footpad of Mxra8-deficient mice, we hypothesized that the CHIKV E1-M88L variant would also be Mxra8-independent in mouse fibroblasts *in vitro*. We infected control NIH-3T3 mouse fibroblast cells, Mxra8-deficient 3T3 cells, or Mxra8-deficient cells expressing Mxra8 *in trans* with each virus and harvested virus containing supernatants at 24 and 48 hours post infection (**Fig. 5D and 5E**). We found that in control 3T3 (**Fig. 5D, red lines**), the E1-M88L variant was attenuated in growth over two days. In addition, the E1-M88L variant replicated slightly worse than WT CHIKV in Mxra8-deficient cells, suggesting that the CHIKV E1-M88L variant did not have an advantage in mouse 3T3 cells. When Mxra8 was introduced *in trans*, we found that WT CHIKV and E1-M88L replication was enhanced, yet the E1-M88L variant was still attenuated (**Fig. 5E, green lines**). These results were confirmed by fixing and staining the cells at the 48 hour timepoint, where we found that that E1-M88L had significantly less infected cells in our control and overexpressed Mxra8 cells (**Fig. 5F**). Finally, we wanted to test whether there was a difference in infectivity of the viruses in 3T3 cells, similar to what we saw in BHK-21 cells (**Fig. 2A**), We incubated each 3T3 cell line with each virus for 1 hour, then replenished the cells with 20 mM ammonium chloride in complete media for 24 hours. After fixing, staining, and quantifying the number of infected cells, we found that again the CHIKV E1-M88L virus was attenuated in both control cells and overexpressed Mxra8 cells (**Fig. 5G**). These findings suggest that while attenuated in our *in vitro* model, E1-M88L is able to infect and replicate within Mxra8-deficient mice *in vivo*, suggesting a cell-type specific mechanism present in mice may be important for virus binding and entry.

### Molecular computational simulations of CHIKV E1 variants reveal changes in E1 and E2 dynamics

To gain mechanistic insight into how the CHIKV E1-M88L variant may be contributing to virus entry, we took a molecular dynamics approach to understand how changes in E1 may be influencing the dynamics of the CHIKV E1-E2 heterodimer *in silico*. By performing molecular dynamic simulations of the prefusion conformation of the E1-E2 heterodimer and doing a PCA analysis of the concatenated trajectory (**Fig. 6A**), we observed that the first Principal Component (PC) was associated with the bending of the envelope glycoproteins of CHIKV and the second PC was associated with the opening of E2 domain B, which is present in a stable conformation between open and closed states (**Fig. 6B**). As a proof of principle, we then applied this approach to the CHIKV E1-V80L variant which we characterized to have defects in entry and is attenuated *in vivo* (20). Importantly, we found that many of these phenotypes were rescued experimentally by the second-site mutation E1-V226A. When we introduced the E1-V80L variant into the simulation we observed a shift in both the second PC, with E2 now in more open conformation (**Fig. 6C** **-light green**). However, when we introduced E1-V80L with the E1-V226A variant we found that E2 was restored to the wild-type conformation (**Fig. 6C** **– dark green)**. As the major difference was observed in PC2, which involves mainly residues in E2-B, we performed a PCA of E2 alone. Consistently in the PC1, we observed the E2 WT protein to be in two distinct states consisting of one major closed state and an open sub-state, with a shift to the open conformation in the E1-V80L mutant which was restored to the WT phenotype by the E1-V226A second-site mutation (**Fig. 6C**). This proof-of-principle highlights that we can understand glycoprotein function through molecular simulations coupled with experimentation.

**Figure 6:**
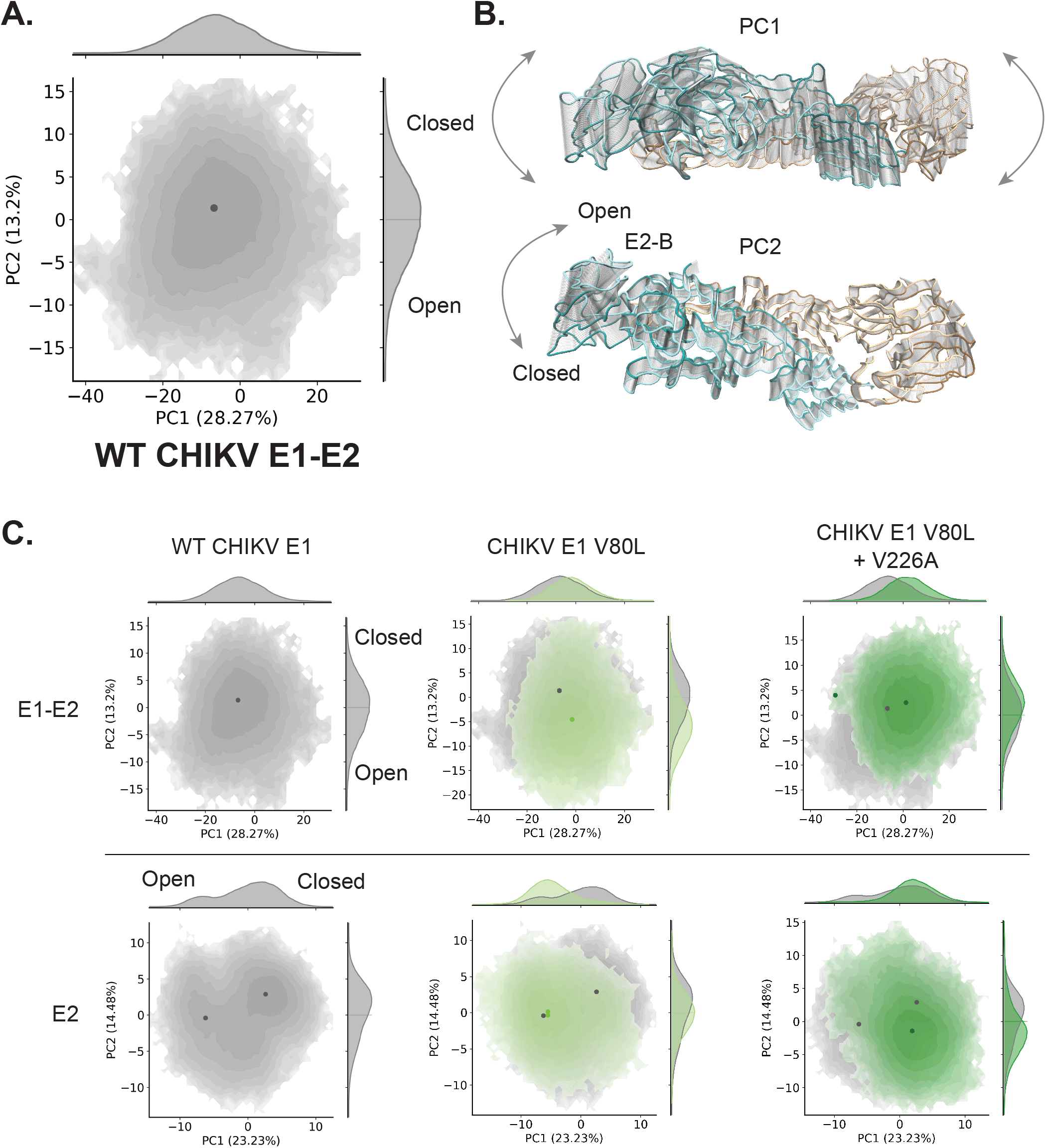
Molecular dynamic simulations of CHIKV E1-M88L variant. (A) Free energy landscape along the first two Principal Components (PC1 and PC2) obtained from a PCA of the Cα atoms of the MD trajectory of the WT E1-E2 heterodimer. A dark dot represents the minimum energy conformation. The explained variance of each PC is shown in the X and Y labels as a percentage of the total variance. (B) Collective motion of the Cα atoms represented by each PC. The extreme conformations are colored in dark or light cyan for E2 and orange for E1 and intermediate conformations are depicted as transparent tubes. (C) Free energy landscape along the first two PCs obtained from the MD trajectory for the WT, E1-V80L or E1-V80L/V226A variants. In the top row is represented the projection along the first two PCs for the E1-E2 heterodimer and on the bottom row the projection only for E2 protein.

When we introduced the E1-M88L variant into the molecular dynamics simulation, we observed that the E2 small open state was largely absent compared to wild-type E2, suggesting that changes in the E1 fusion loop can place the E2 protein into a more closed conformation (**Fig. 7A**). Finally, given that we identified the E1-M88L variant alongside of E1-V80Q in mosquitoes, we asked whether the E1-V80Q:E1-M88L double variant changed the simulation of E1-V80Q. When modeled alone, we found that the E1-V80Q variant shifted E2 more to the closed conformation and E1-M88L further shifted E2 to this closed state and did not restore the wild-type phenotype (**Fig. 7A**). These results suggest that changes in E1 can significantly change the dynamics of E2. To confirm these findings, we ran simulations on several other E1-V80 variants we characterized in previous studies to increase virulence and transmission (E1-V80I) or genetically unstable (E1-V80K, V80F, and V80E) (**Fig. 7B**). Although each of these variants significantly changed the dynamics of E2 in different ways, there seems to be an association of a shift to the open state of E2 domain B with a reduced phenotype (E1-V80L, V80K, V80F and V80E). Taken together, these molecular simulations highlight dramatic changes in E2 dynamics imparted by E1 variants and provide molecular insight into how E1 can drive cell binding and infectivity in multiple ways.

**Figure 7:**
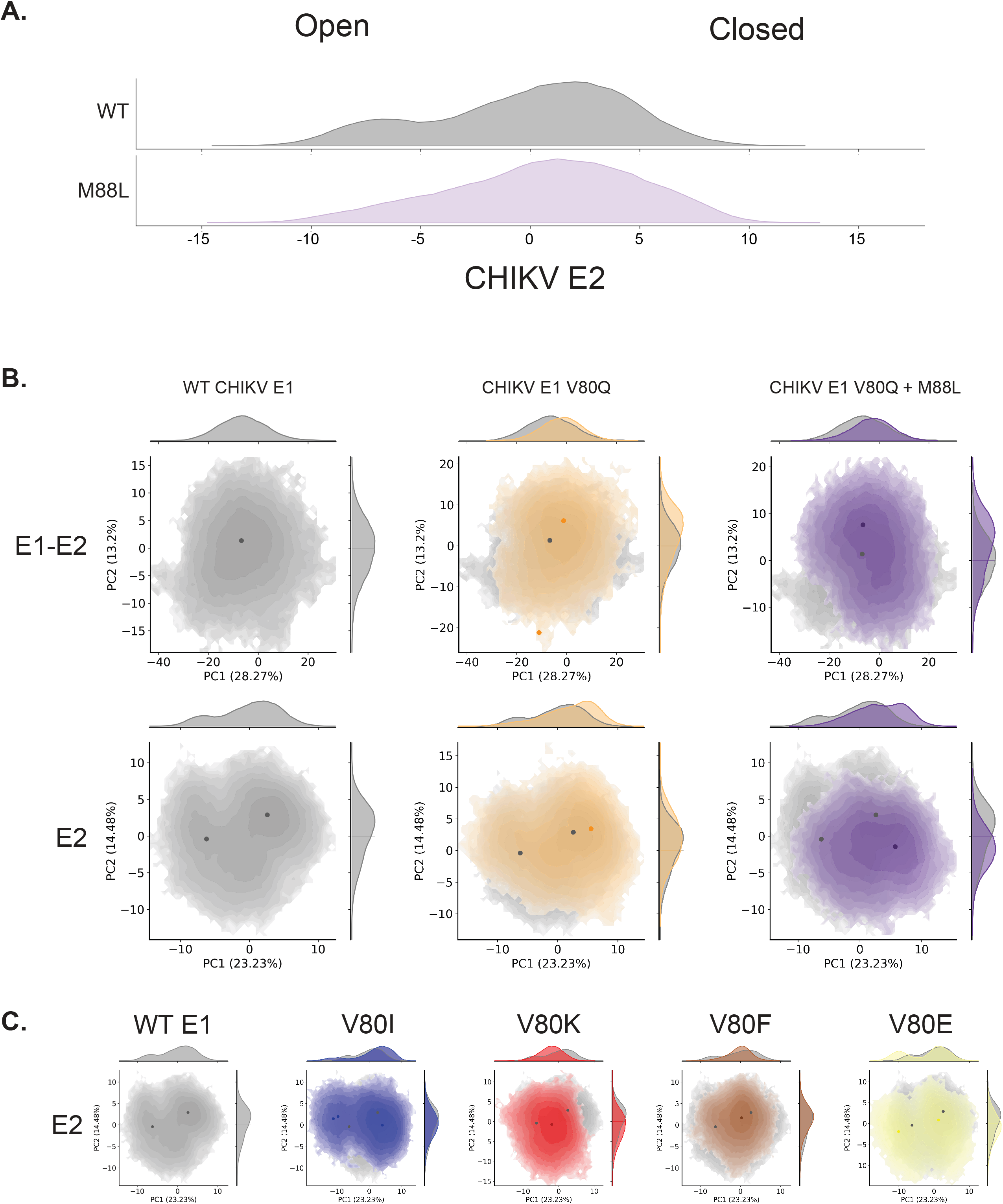
CHIKV E1 variants influence E2 molecular dynamics *in silico*. (A) Free energy landscape along the first PC obtained for E2 protein of WT and E1-M88L variant. (B) Free energy landscape along the first two PCs obtained from a PCA of the Cα of the MD trajectory for E1-E2 heterodimer (top) or E2 protein (bottom). The projection along PC1 and PC2 is shown as in figure 6A for WT, E1-V80Q and E1-V80Q/M88L variants. (C) Free energy landscape along the first two PCs obtained for E2 protein with WT, E1-V80I, E1-V80K, E1-V80F and E1-V80E variants.

## Discussion

Chikungunya virus (CHIKV) is an alphavirus that has caused significant outbreaks worldwide, including recent explosive outbreaks in South America (9) (10). Our previous studies using CHIKV as a model alphavirus have examined how the E1 class II fusion glycoprotein contributes to viral fusion, infectivity and evolution (20) (24) (25). We identified two second-site E1 glycoprotein variants (E1-M88L and E1-N20Y) in the bodies of *Ae*. *aegypti* mosquitoes infected with an attenuated E1 variant, CHIKV E1-V80Q, that corresponded with an increased plaque size phenotype (20). This initial observation allowed us to hypothesize that the emergence of E1-M88L and/or E1-N20Y was able to rescue the attenuation of CHIKV E1-V80Q. Unfortunately, due to the genetic instability of E1-V80Q, we were unable to answer this specific question, and instead used *in vitro* and *in vivo* approaches to understand how E1-M88 and E1-N20 contribute to the CHIKV life cycle.

When we focus on the location and conservation of E1-N20 and E1-M88 across different alphaviruses, we find that E1-N20 in the E1 domain I hinge is the most variable of the two with the majority of alphaviruses containing a bulky amino acid at this position. The domain I hinge has been shown to be important for virus entry and for domain swiveling during membrane fusion (26) (27). Given that the E1-N20Y variant increases infection in C6/36 cells and not mammalian cells, it could suggest that this residue is important for insect-specificity between alphaviruses for entry. In contrast, residue E1-M88, in the fusion loop, is highly conserved amongst alphaviruses, and is flanked by a conserved phenylalanine that has been shown to be critical for infectivity in multiple alphaviruses (28). Interestingly AURV, UNAV, and MIDV all encode a leucine at this position, suggesting there may be an evolutionary role for the residue in alphaviruses.

Using *in vitro* approaches, we observed minimal differences in non-structural and structural protein accumulation, RNA replication, and growth kinetics of E1-M88L and E1-N20Y in BHK-21 and C6/36 cells compared to wild-type CHIKV. However, we did find that E1-N20Y enhanced infection specifically in C6/36 cells while E1-M88L was able to enhance binding and infectivity in BHK-21 and C6/36 cells. These results suggest that the fusion loop and the hinge region may contribute to the cell- and/or host-specific entry of CHIKV. Interestingly, previous work has shown that Semliki Forest virus E1-M88L had minimal effects on membrane fusion, surface expression, antibody binding and glycosylation (29). These results suggest that the function of residue E1-M88 may be virus-specific or study-specific depending on lab cell lines and culture conditions. Nonetheless, the idea of cell- and/or host-specificity is supported by experiments in NIH-3T3 cells, where we found that E1-M88L was attenuated in these cells, suggesting that there are host and/or cell-dependent differences. This hypothesis may also explain why we observe no differences in growth kinetics in BHK-21 and C6/36 cells, even with enhanced infectivity of the variants. These results may be due to the fact that the growth curves were tittered on Vero cells, and variations in Vero-specific phenotypes of these variants may contribute to differences in reported growth kinetics as compared to viral infectivity. Future work focusing on how the CHIKV E1 fusion loop and hinge impact infectivity of multiple cell types may address these questions and shed light on cell- and host-dependent entry.

Given our enhanced infection of BHK-21 and C6/36 cells with E1-M88L, we also addressed whether CHIKV E1-M88L had an advantage in mosquitoes and mice. We found no major differences between E1-M88L and wild-type CHIKV in mosquitoes or mice, with the exception of a decrease in viral titers in footpad and increased clearance from the serum at 3 days post infection. One explanation of these findings may be that, as we saw with *in vitro* NIH 3T3 experiments, E1-M88 is critical for cell and/or host dependent entry and therefore results on BHK-21 cells do not reflect what happens in mice. In addition, C6/36 cells are *Aedes albopictus* cells and we infected *Ae. aegypti* mosquitoes, suggesting there could be species differences as well. One interesting observation was that CHIKV E1-M88L lead to increased viral titers in the footpad of Mxra8-deficient mice, suggesting that E1-M88L can facilitate entry in the absence of a receptor. These results are in line with the increased cell binding and infectivity we observed *in vitro* and while E1-M88L was dependent on Mxra8 and attenuated in NIH 3T3 cells *in vitro*, the complex *in vivo* environment could suggest that E1-M88L is able to better infect other cell types or bind to other host molecules for entry. Future studies exploring the broad cell tropism of CHIKV E1-M88L in mammals and mosquitoes will be critical to mapping out how this residue in the fusion loop contributes to entry.

Finally, given that CHIKV E2 is the attachment protein needed for interactions with Mxra8 and glycosaminoglycans (GAGs) (2), we found it interesting that changes in the E1 fusion loop, would impact binding so dramatically. Using molecular dynamic simulations, we observed that modeling changes in CHIKV E1 domain II, the fusion loop and β-strand c, led to changes in not only E1 but also E2 dynamics, potentially explaining what we see experimentally. Indeed, we have observed changes in cell binding and GAG interactions with CHIKV E1 variants in the E1-E1 interface (24), supporting the idea that changes in E1 can have multiple contributes to glycoprotein structure and dynamics. It will be important to better understand how E1 and E2 work together using complementary *in silico* and laboratory experiments to better understand the mechanisms of CHIKV entry.

Taken together, alphaviruses are a diverse genus that exhibits high epidemic potential due to their prevalence and transmission globally. These viruses are a leading cause of arthritic and encephalitic disease, yet there are limited antiviral therapeutics available. Therefore, understanding the molecular mechanisms of how alphaviruses infect both insects and mammals, as well as specific cell types during infection is essential to our understanding of viral emergence and the development of antiviral therapies.

## Acknowledgements

We thank all members of the Stapleford Lab for helpful discussion on this project. We thank Dr. Meike Dittmann at the NYU School of Medicine for use of the CX7 Cell-Insight microscope and Drs. Ludo Desvignes and Dominick Papandrea for use of the NYU Grossman School of Medicine ABSL3 facility. This work was supported by funding from the NYUGSoM Start up, the American Heart Association Postdoctoral Fellowship (19-A0-00-1003686) (M.G.N), NIH/NIAID R01 AI162774-01A1 (K.A.S), PICT 2020-3371 (D.E.A.).

## Materials and Methods

### Cell lines

Baby hamster kidney cells (BHK-21, ATCC CCL-10) and NIH-3T3 cells (gifts from Dr. Michael Diamond at Washington University (22)) were grown in Dulbecco’s Modified Eagle Medium (DMEM, Corning) with 10% fetal bovine serum (FBS; Atlanta Biologicals), 1% nonessential amino acids (NEAA, Fisher Scientific) and 10 mM HEPES (Invitrogen). Vero cells (ATCC CCL-81) were grown in DMEM with 10% newborn calf serum (NBCS, Gibco). Mammalian cells were maintained at 37°C with 5% CO_2_. *Aedes albopictus* cells (C6/36, ATCC CRL-1660) were grown in L-15 media (Corning) supplemented with 10% FBS, 1% NEAA and 1% tryptose phosphate broth (Invitrogen), and were maintained at 28°C with 5% CO_2_. All cell lines were confirmed mycoplasma free.

### Biosafety

All work with chikungunya virus (CHIKV) and CHIKV E1 variants were completed under Biosafety level 3 (BSL3) conditions at the NYU Grossman School of Medicine. Mouse and mosquito work were completed in the NYU Grossman School of Medicine ABSL3 animal and insect facility. Animal work was complaint with NYU Grossman School of Medicine Institutional Animal Care and Use Committee (IACUC) protocol #IA16-01783.

### Viruses

Wild-type CHIKV (strain 06-049), E1 glycoprotein variants, and viral derivatives expressing ZsGreen or Firefly luciferase reporter proteins under the subgenomic promoter were produced from infectious clones as previously described (30). E1 glycoprotein variants were introduced in the CHIKV infectious clone by site-directed mutagenesis using the following primers: E1-M88L; Forward GTCTACCCATTTTTGTGGGGCGGCG, Reverse CGCCGCCCCACAAAAATGGGTAGAC, E1-N20Y; Forward GTATAAGACTCTAGTCTATAGACCTGGCTACAG, Reverse CTGTAGCCAGGTCTATAGACTAGAGTCTTATAC. The E1 variants were introduced into an infectious clone expressing ZsGreen or Firefly luciferase by subcloning the XhoI/NotI restriction fragment from the unmarked CHIKV clone into the same restriction sites of the reporter plasmids. All plasmids were Sanger sequenced to confirm the E1 variants and that there were no second-site mutations.

To generate infectious virus, 10 μg of each infectious clone plasmid was linearized overnight with NotI restriction endonuclease (Invitrogen), purified by phenol:chloroform extraction and ethanol precipitation, and resuspended in nuclease free water (20) (30). CHIKV RNA was *in vitro* transcribed using the SP6 mMessage mMachine kit (Invitrogen) following the manufacturer’s instructions. RNA was purified by phenol:chloroform extraction and ethanol precipitation, diluted to 1 μg/μl in nuclease free water, aliquoted, and stored at -80°C. BHK-21 cells (10^7^ cells/ml) were electroporated with 10 μg of *in vitro* transcribed RNA by 1 pulse of 1,200 V, 25 Ω, and infinite resistance. Electroporated cells were added to a T25 flask in complete media (DMEM, 10% FBS, 1% NEAA and 1% HEPES). After incubation at 37°C for 72 hours, the passage 0 (p0) supernatant was centrifuged at 1,200 rpm for 5 minutes and was used to infect a T-175 flask of BHK-21 cells to produce a passage 1 (p1) working stock. After incubation at 37°C for 24 hours, p1 virus was centrifuged at 1,200 rpm for 5 minutes, aliquoted and stored at -80°C. Viral RNA was extracted and all viruses were Sanger sequenced to address genetic stability as described below. Infectious virus titers were quantified by plaque assay for all stocks as described below. Ultracentrifugation was used to generate purified virus stocks. Viruses were pelleted over a 20% sucrose cushion by centrifugation at 25,000 rpm for 4 hours. Purified virus particles were resuspended in media consisting of DMEM containing 2% FBS. Viral titers were quantified by plaque assay and viral genomes were quantified by RT-qPCR as described below.

### Plaque assay

Infectious virus was quantified via plaque assay. Supernatants containing virus were diluted 10-fold in DMEM and added to a monolayer of Vero CCL81 cells (350,000 cells/well) in a 12-well plate. After a 1 hour incubation at 37°C, media comprised of DMEM, 2% FBS, and 0.8% agarose was added to each well and incubated at 37°C for 72 hours. For all mosquito and mouse samples, 1% antibiotic-antimycotic (Gibco) was added to the media. Cells were fixed with 4% formalin for 1 hour, agarose plugs removed, and plaques were visualized using crystal violet. Viral titers were calculated using the lowest countable dilution.

### RNA extractions and RT-qPCR

CHIKV RNA for qPCR assays was extracted and purified using TRIzol reagent (Fisher Scientific) following the manufacturer’s protocol. To quantify the number of viral RNA genomes/mL, a TaqMan RNA-to-CT one-step kit (Applied Biosystems, Fisher Scientific) was used with the following primers targeting the nsP4 region of the genome: Forward 5’TCACTCCCTGCTGGACTTGATAGA 3’. Reverse

5’TTGACGAACAGAGTTAGGAACATACC 3’. Probe 6919-FAM

5’AGGTACGCGCTTCAAGTTCGGCG 3’. A standard curve was generated from 10-fold dilutions of *in vitro* transcribed RNA was included for all samples and all were run in technical duplicate.

### Sanger Sequencing

Viral RNA was extracted from stocks as described above. cDNA was generated using a Maxima H Minus Enzyme First Strand cDNA Synthesis Kit (Thermo Scientific) following the manufacturer’s instructions. The CHIKV genome was amplified in 5 PCR fragments using Phusion High-Fidelity PCR Master Mix Kit with HF Buffer (Thermo Scientific). CHIKV fragments were confirmed by an agarose gel and sent to Sanger Sequencing using the primers in Table 1 for confirmation of mutations in all working CHIKV viral stock mutations (Genewiz).

**Table 1:**
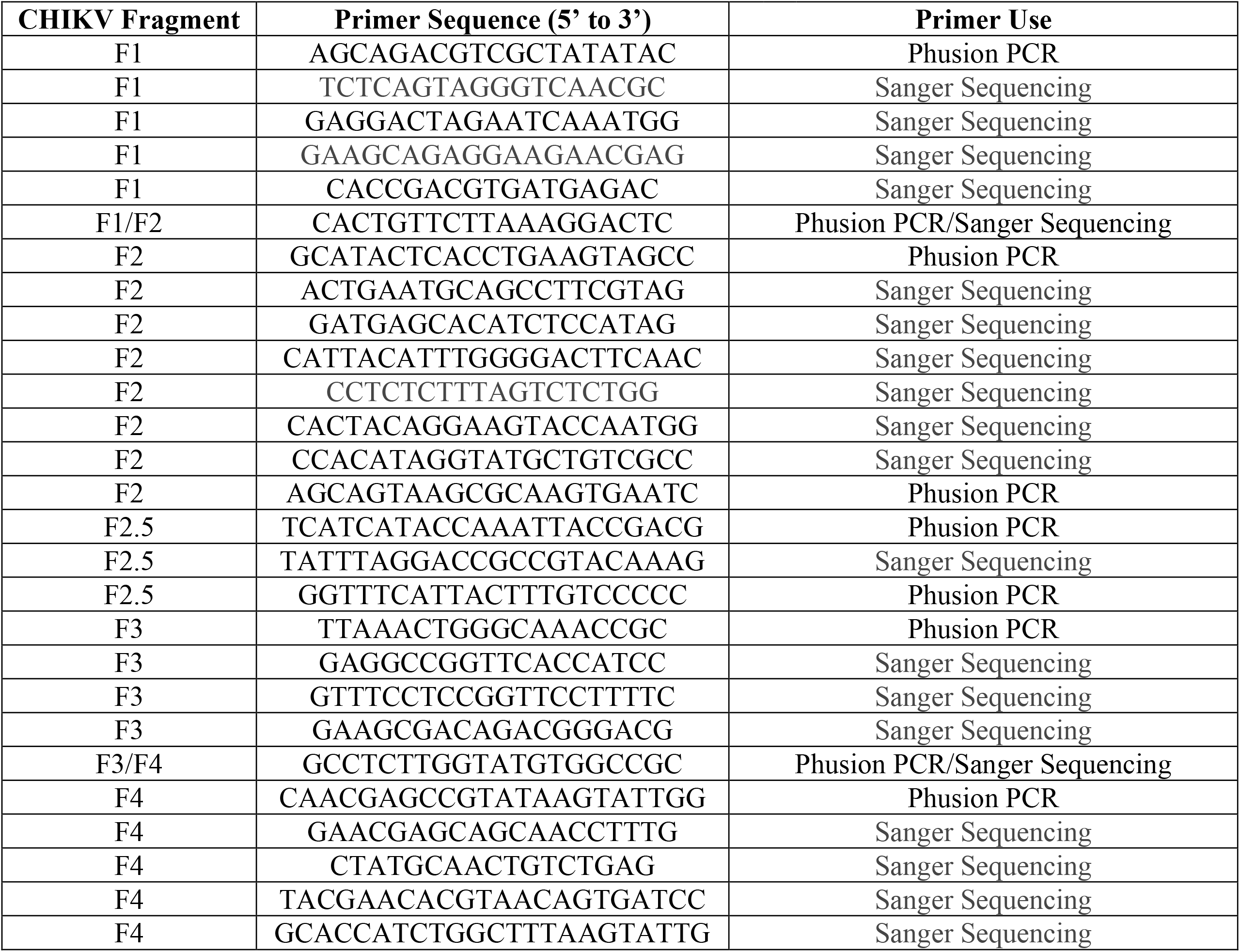
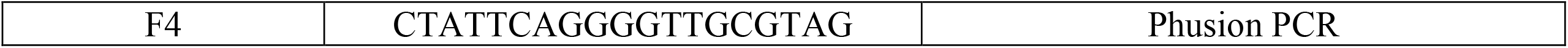
PCR and sequencing primers used in this study.

### Cell-Insight CX7 quantification of infected cells

Cells infected with CHIKV-ZsGreen virus were fixed with 4% paraformaldehyde (PFA) for 1 hour at room temperature (20). Following incubation, fixed cells were stained with 4’,6-diamidino-2-phenylindole (DAPI, Thermo Scientific) diluted 1:1000 in phosphate-buffered Saline (PBS, Corning) for 1 hour at room temperature. A Cell-Insight CX7 high-content microscope (Thermo-Scientific) was used to quantify the percent of infected cells in each well, by dividing the number of ZsGreen positive cells with the total number of DAPI stained cells.

### Western blotting

BHK-21 cells were electroporated with 10 μg of *in vitro* transcribed RNA and seeded in a T-25 flask for 48 hours at 37°C. After incubation, cells were washed once with PBS and scraped off the flask into cold PBS. Cells were pelleted by centrifugation at 1,200 rpm for 5 mins at 4°C and resuspended in lysis buffer (1X Tris buffered saline (TBS; 20 mM Tris-HCl, pH 7.4, 150 mM NaCl, 1 mM EDTA, 1% TX-100 and 1X Halt protease inhibitor cocktail (Thermo) for 1 hour on ice. After lysis, debris was removed by centrifugation at 10,000 x g for 5 mins and mixed 1:1 with 2X Laemmli buffer containing 10% β-mercaptoethanol. Samples were boiled at 95°C for 10 minutes and centrifuged at 10,000 x g for 5 mins. Proteins were separated by SDS-PAGE on a Mini-PROTEAN TGX Stain-Free pre-cast polyacrylamide gel (Bio-Rad), transferred to a hydrophobic polyvinylidene fluoride (PVDF) membrane (Immobilon), and incubated in blocking buffer comprised of 1X TBS-T (TBS + 0.1% Tween-20) and 5% dry milk overnight at 4°C. Blots were then incubated with primary CHIKV antibodies including anti-rabbit CHIKV-E1 (gift from Dr. Gorben Piljman), anti-mouse CHIKV-E2 (CHIK-48, BEI Resources), anti-rabbit CHIKV-Capsid (gift from Dr. Andres Merits), anti-rabbit CHIKV-nsP1 (gift from Dr. Andres Merits) and anti-mouse Actin (MA5-11869, ThermoFisher) for 2 hours at room temperature. After multiple washes with 1X TBST, blots were incubated with anti-rabbit or anti-mouse IgG HRP secondary antibodies (Invitrogen) for 1 hour at room temperature. After another round of washes, blots were developed using a SuperSignal West Pico plus chemiluminescence substrate kit (Thermo) and imaged using a ChemiDoc MP imaging system (Bio-Rad). Images were analyzed using Image Lab (Bio-Rad).

### Growth curves

BHK-21 cells (50,000 cells/well) or C6/36 cells (200,000 cells/well) were seeded in a 24-well plate 24 hours prior to infection. Cells were infected with each virus diluted in DMEM at an MOI of 0.1 for 1 hour at 37°C (BHK-21) or 28°C (C6/36). After incubation, cells were washed twice with PBS and complete BHK media or insect media was added to the cells. Supernatants were collected from each corresponding timepoint and infectious virus was quantified via plaque assay as described above.

### Ammonium chloride bypass assay

BHK-21 cells (15,000 cells/well) were seeded in a Costar 96-well plate and incubated at 37°C 24 hours prior to infection. After the seeded plate was incubated at 4°C for 1 hour, ZsGreen expressing viruses were added to the cells at an MOI of 1 and incubated at 4°C for another hour. A final concentration of 20 mM ammonium chloride (NH_4_Cl) was diluted in complete media, then added to each well at its corresponding timepoint and incubated overnight at 37°C. The following day, infected cells were fixed with 4% PFA and quantified by the CX7 as stated above.

### C6/36 Infectivity Assay

C6/36 cells (100,000 cells/well) were seeded in a 96-well plate at 28°C for 24 hours. Cells were then infected with WT CHIKV or an E1 mutant virus at an MOI of 0.1 and incubated at 28°C for 1 hour. A final concentration of 20 mM ammonium chloride was diluted in insect cell media and treated on the cells for incubation at 28°C. After 24 hours, cells were fixed with 4% PFA, stained with DAPI, and quantified using a CX7 high-content microscope.

### Virus binding assay

BHK-21 cells (100,000 cells/well) or C6/36 cells (350,000 cells/well) were seeded in a 12-well plate and incubated at 37°C or 28°C the day before infection. Cells were pre-treated with binding buffer media compromised of DMEM, 0.2% bovine serum albumin (BSA), 10 mM HEPES and 20 mM NH_4_Cl at 4°C for 1 hour. After incubation, cells were washed once with PBS, then incubated with purified virus at an MOI of 100 (based on RNA genomes) diluted in binding buffer and left for 30 mins on ice. Virus media was removed and cells were washed with cold PBS three times. Cells were then collected in Trizol and RNA was extracted to quantify viral RNA genomes via RT-qPCR as described above.

### Cholesterol-depletion assay

BHK-21 cells (20,000 cells/well) were seeded in a 96-well plate and incubated for 24 hours at 37°C. Cells were pretreated with increasing concentrations of methyl-β-cyclodextrin (MβCD) for 1 hour at 37°C. Following the incubation, the cells were washed once with PBS, and then infected with each corresponding CHIKV-ZsGreen virus at an MOI of 1 at 37°C for 1 hour. A final concentration of 20 mM ammonium chloride was added to the cells and were then incubated at 37°C for 24 hours. Infected cells were fixed with 4% PFA, stained with DAPI and quantified using a CX7 high-content microscope.

### Luciferase assay

BHK-21 (20,000 cells/well) were seeded in a 96-well plate 1 day prior to transfection. Cells were transfected with 10 mg of WT CHIKV, CHIKV nsP4-GNN or E1-M88L RNA using Lipofectamine 2000. Lipofectamine was mixed with Opti-MEM (Thermo Scientific) and incubated at room temperature for 10 minutes. This was then added to a mixture of RNA and Opti-MEM at a at 1:2 ratio of Lipofectamine to RNA. After incubating for 5 minutes at room temperature, 10 µl of this mix was added to cells and was incubated at 37°C. At 4, 6, 8 and 24 hours post-transfection, supernatants were collected and cells were lysed with Steady-Glo firefly buffer reagent (Pierce) in a white 96-well plate. Collected supernatants were used to infect BHK-21 cells (20,000 cells/well) for 24 hours, and luciferase activity was read as before.

Luminescence was read using a SpectraMax M3 plate reader (Molecular Devices), and raw data was quantified as relative luciferase units.

### NIH 3T3 virus growth and infectivity assays

Control NIH-3T3 cells (Control + Vector), 3T3 *ΔMxra8* receptor knockout cells (ΔMxra8 + Vector) and 3T3 *ΔMxra8* cells overexpressing *Mxra8 (ΔMxra8 + Mxra8*) *in trans* were seeded as 20,000 cells/well in a 96-well plate 1 day prior to infection (22). Cells were infected with a purified WT CHIKV-ZsGreen or E1-M88L-ZsGreen at an MOI of 5 and incubated at 37°C for 1 hour. Cells were then either given 20 mM ammonium chloride with complete media and fixed after 24 hours post-infection (hpi), or were given complete media and fixed at 48 hpi.

Supernatants were collected at both 24 and 48 hpi from both experiments, and were used for a plaque assay to determine infectious particles. Once cells were fixed, they were stained with DAPI for 30 minutes, then imaged and quantified using a Cell-Insight CX7 high-content microscope.

### Mouse infections

Wild-type C57BL/6J (Jackson Laboratory) and *Mxra8^Δ8/Δ8^* mice (a gift from Dr. Michael Diamond (21)) were breed and maintained in the NYU Grossman School of Medicine BSL2 vivarium and moved to the ABSL3 facility prior to and during infection. *Mxra8^Δ8/Δ8^* mice were rederived from sperm at the NYU Grossman School of Medicine. WT and *ΔMxra8* littermate controls were generated by first crossing *Mxra8^Δ8/Δ8^* mice to WT C57BL/6J mice to generate heterozygote *Mxra8^+/Δ8^* mice and subsequently crossing these heterozygotes to obtain WT and *Mxra8^Δ8/Δ8^* mouse lines. All mice were genotyped (Transnetyx) prior to each experiment.

For virus infections, male and female 5–7-week-old mice were anesthetized using isoflurane and infected with 1000 PFU of WT CHIKV or the E1-M88L variant via footpad injection. At each corresponding timepoint, mice were euthanized via CO_2_ inhalation. Serum, ipsilateral footpad, and ipsilateral muscle from each mouse was harvested and placed in separate 2 mL tubes containing 500 μl PBS and two 5 mm stainless steel balls. Samples were homogenized using a Tissue-Lyser II (Qiagen) for 2 minutes at 30 Hz, centrifuged at 8,000 rpm for 8 minutes, then used to quantify infectious particles via plaque assay.

### Mosquito infections

*Aedes aegypti* (Poza Rica, Mexico) mosquitoes were reared and maintained in a 28°C incubator with 70% humidity in an ABSL3 insectary (31) (24). Mosquitoes 4-7 days post-emergence were infected with virus via blood meal through a pig intestine membrane. Each virus was mixed with PBS-washed defibrinated sheep blood (Fisher Scientific) containing 5 mM ATP to a concentration of 10^6^ PFU/mL and fed to ∼50-60 female mosquitoes. After 1 hour, engorged mosquitoes were sorted and maintained for 7 days, after which their bodies, legs and wings were separated into 2 mL tubes containing 200 μl PBS and one 5 mm stainless steel balls. Samples were then homogenized and ground using a Tissue-Lyser II (Qiagen) for 2 minutes at 30 Hz, centrifuged at 8,000 rpm for 8 minutes, then used to quantify infectious particles via plaque assay.

### Molecular Dynamic simulations

To study the impact of E1-M88L mutation in the envelope glycoproteins function at a molecular level, we performed unbiased atomistic Molecular Dynamics simulations in explicit solvent. As an initial structure, we used the pre-fusion conformation of the E1-E2 heterodimer (PDB: 3N42). To simulate the variants, we introduced the point mutations in the WT protein using PyMol (Schrödinger L, DeLano W. PyMOL. In: http://www.pymol.org/pymol). All simulations were done using the GROMACS 2021.2 software package (https://doi.org/10.1016/j.softx.2015.06.001). For each variant, the protonation state at pH 7 of all protonable residues was determined using PROPKA [https://doi.org/10.1021/ct100578z]. The Amber99SB*-ILDN force field [https://doi.org/10.1002/prot.22711, https://doi.org/10.1021/jp901540t] was employed to describe the protein. A cutoff value of 10 Å was applied for short-range electrostatic and van der Waals interactions. Long-range electrostatic interactions were treated with PME (Particle Mesh Ewald). The system was solvated with a dodecahedral box of TIP3P water [https://doi.org/10.1063/1.445869]. The water box extended 12 Å from the protein surface and the system was neutralized with 0.15 M NaCl. The system was minimized using the Steepest Descent algorithm. Subsequently, it was heated to 310 K and equilibrated for 200 ps in the NVT ensemble using the V-rescale thermostat [https://doi.org/10.1063/1.2408420]. A second equilibration step was performed for 1 ns in the NPT ensemble using the Berendsen barostat [https://doi.org/10.1063/1.448118]. Finally, a production run was conducted in the NPT ensemble using the V-rescale thermostat and the Parrinello-Rahman barostat [https://doi.org/10.1063/1.328693]. In both equilibration and production runs, the LINCS algorithm [https://doi.org/10.1002/(SICI)1096-987X(199709)18:12<1463::AID-JCC4>3.0.CO;2-H] was used and a time step of 2 fs was employed. For each variant we performed two independent runs of 250 ns starting from the same minimized structure. All simulations were performed on the high-performance computing centers of NYU Langone and CCAD (https://ccad.unc.edu.ar/).

We analyzed the final 200 ns of each MD run. We concatenated the trajectory of both replicas of all variants and we performed a PCA over the concatenated trajectory. In order to compare the conformational space sampled by each variant, we projected the trajectory of each one on PC1 and PC2. Using the simulation done with E1-E2 heterodimer we performed the PCA over both E1 and E2 or over E2 alone. The preprocessing of the trajectories and the PCA analysis were done using GROMACS tools. The 2D projections and graphics were done using an in-house Python script using Pandas, Scipy, Seaborn and Matplotlib packages.

### Protein structures and sequencing analysis

The CHIKV E1 and E2 glycoprotein (PDB: 3N42) was rendered in PyMol (Version 2.5.2). CHIKV nucleotide sequencing alignments were generated with SeqMan Ultra (DNA Star), and alphavirus amino acid alignments were generated with MegAlign Pro (DNA Star). The following alphavirus were used for E1 amino acid alignments: chikungunya virus (06-049), Mayaro virus (NP_740694), Sindbis virus (AWT57845), Ross River virus (QTC33398), Eilat virus (YP_006732328), Semliki forest virus (CAA52444), Middelburg virus (AAL35777), O’nyong n’yong virus (YP_010775618), Aura virus (AWQ38331), Una virus (AAL35783), western equine encephalitis virus (QEX51909), eastern equine encephalitis virus (AAU95735).

### Statistics and data analysis

All data were analyzed using GraphPad Prism (Version 9). All *in vitro* experiments were completed with at least two biological replicates and internal technical duplicates (exact details can be found in the figure legends). All *in vivo* experiments were completed with at least n=6 for mice and n=43 for mosquito infections. P value of < 0.05 is considered statistically significant.

